# Full Mitochondrial Phylogeny of Cutthroat Trout with Comments on Species Delimitation and Taxonomy

**DOI:** 10.1101/2025.09.30.679330

**Authors:** Tanner S. Van Orden, Kevin B. Rogers, Peter Searle, Andrea L. Kokkonen, Dennis K. Shiozawa, R. Paul Evans

**Affiliations:** Department of Biology, Brigham Young University, Provo UT, USA; Aquatic Research Section, Colorado Parks and Wildlife, PO Box 775777, Steamboat Springs, Colorado 80477, USA; Department of Ecology and Evolutionary Biology, Cornell University, Ithaca, NY, USA; Department of Microbiology and Molecular Biology, Brigham Young University, Provo UT, USA

## Abstract

Despite the important role that species concepts and taxonomy play in understanding the evolution of organisms, taxonomists have struggled to describe the relationship between distinct lineages of Cutthroat Trout. This struggle inspired a special workshop at the 2015 annual meeting of the American Fisheries Society that emphasized the need for a revised taxonomy of Cutthroat Trout. To further assess the relationship between different Cutthroat Trout lineages, we sequenced and assembled full mitochondrial genomes from 123 Cutthroat Trout from across their native range. We used maximum likelihood and Bayesian phylogenetic approaches to examine the evolutionary relationships between all named Cutthroat Trout subspecies. Through these analyses, we find 11 lineages of Cutthroat Trout that diverged > 2 million years ago, and at least 13 lineages that diverged > 1 million years ago, highlighting the ancient origin of Cutthroat Trout diversity. Despite the ancient split between many Cutthroat Trout lineages, we do not find that Coastal, Westslope, Lahontan, or Yellowstone Cutthroat Trout are more distinct than other lineages, and as such, we continue to recognize a single Cutthroat Trout species (*Oncorhynchus clarkii*) comprised of twelve distinct subspecies, maintaining consistency with conservation efforts, management plans, ESA-related decisions, while also preserving stability associated with historic nomenclature.

## Introduction

Species concepts and taxonomy have always been an essential part of understanding the evolution of organisms (Mayr, 1968). Additionally, species concepts play an important role in the protection and conservation of species, yet what defines a species remains elusive. Over the last 80 years, at least 34 concepts have been proposed, but no consensus on which species concept is the most accurate or useful has been achieved (Frankham et al., 2012; Mayden, 1999; Mayr, 1968; Zachos, 2018). While new concepts continue to proliferate (Hill, 2017; Shanker et al., 2017; Zachos, 2018), most tend to drift from Mayr’s biological species concept, which was the accepted standard for decades. Instead of using reproductive isolation to delineate species, these newer approaches acknowledge even finer diversity. For example, the phylogenetic species concept allows any monophyletic group to be treated as a species, while some practitioners have used the Unified Species Concept to justify splitting metapopulations into separate species (De Queiroz, 2007; Nixon & Wheeler, 1990). While this transition helps highlight complex taxonomic relationships, it has necessarily resulted in the rapid inflation of numbers of described species within well studied taxonomic groups and shaken stability in nomenclature – a stated objective of the International Commission on Zoological Nomenclature and the American Fisheries Society (Page et al. 2023).

Cutthroat Trout (*Oncorhynchus clarkii*) have been classified into many different taxonomic groupings following their discovery by the Western world in 1541, particularly in the 1800s (Behnke, 1992, 2002; Trotter, 2008). Taxonomists have struggled to describe the diversity found within Cutthroat Trout, with as many as 40 separate species being recognized before many were synonymized as either subspecies of Rainbow Trout (*Salmo gairdnerii*) or Cutthroat Trout (*Salmo clarkii*) (Jordan et al., 1930; Miller, 1950). In 1989, Rainbow Trout and Cutthroat Trout were moved into the genus *Oncorhynchus* to reflect the close genetic relationship that they share with Pacific salmon (Smith & Stearley, 1989). Cutthroat Trout are native to the Pacific Coast of North America from Alaska south to California, and inland through the Columbia and upper Missouri river basins, the Great Basin (including the Lahontan and Bonneville basins), and along with the headwaters of the Colorado, Rio Grande, South Platte, and Arkansas river basins. Various authors have recognized between 11 and 16 different subspecies of Cutthroat Trout within this native range (Behnke 1979, 2002; Smith & Stearley, 1989; Houston et al. 2012; Penaluna et al. 2016; Budy et al. 2019; Kokonnen et al. 2024).

More recent studies revealed inaccuracies in the taxonomy of Cutthroat Trout at the subspecies level (Bestgen et al., 2019; Loxterman & Keeley, 2012; Metcalf et al., 2007, Metcalf et al., 2012; Shiozawa & Williams, 1992). This precipitated a special workshop at the 2015 annual meeting of the American Fisheries Society that emphasized the need for a revised taxonomy of Cutthroat Trout. Workshop participants settled on the Unified Species Concept as a way to delineate diversity in Cutthroat Trout, as relationships between them are continuously resolved. This workshop aimed to identify lineages across all subspecies that were on separate evolutionary trajectories, which they referred to as uniquely identifiable evolutionary units (UIEUs). After much deliberation, at least 25 UIEUs were recognized across the range of Cutthroat Trout. In November 2023, the Joint Committee of the American Fisheries Society and the American Society of Ichthyologists and Herpetologists moved to reclassify Cutthroat Trout into four species without any named subspecies. These four species include Coastal Cutthroat Trout (*Oncorhynchus clarkii*), Lahontan Cutthroat Trout (*Oncorhynchus henshawii*), Westslope Cutthroat Trout (*Oncorhynchus lewisi*), and Rocky Mountain Cutthroat Trout (*Oncorhynchus virginalis*) (Page et al., 2023). This reclassification was based on an assessment by Markle (2018) who proposed an “interim classification” of Cutthroat Trout before more work could be done to allow a systematic reclassification.

While this interim classification does not alter existing conservation plans for previously recognized Cutthroat Trout subspecies, it did not adequately reflect the diversity described in that same workshop (Campbell et al., 2018; Rogers et al., 2018a; Shiozawa et al., 2018), or from other taxonomic studies (Bestgen et al., 2019; Houston et al., 2012; Loxterman & Keeley, 2012; Metcalf et al., 2007, Metcalf et al., 2012; Pritchard et al., 2009) that reveal substantial genetic diversity between other subspecies on par with that seen between the four new “species”. To clarify these relationships further, we sequenced and assembled full mitochondrial genomes from 123 Cutthroat Trout from across their native range. An additional 25 publicly available mitochondrial genomes across Cutthroat Trout, Rainbow Trout (*Oncorhynchus mykiss)*, and Chinook Salmon (*Oncorhynchus tshawytscha)* were included in all of our analyses. We used maximum likelihood and Bayesian phylogenetic approaches to examine evolutionary relationships across the entire Cutthroat Trout species complex.

## Methods

### Mitochondrial genome acquisition

Samples used in the project were collected by the authors or state and federal fisheries biologists between 1990 and 2022. Isolated DNA and fin clips were housed at the Monte L. Bean Museum (MLBM; Appendix A), or at Pisces Molecular (PM; Boulder, Colorado; Appendix A). DNA sequencing data for this project was generated across multiple independent projects at both Brigham Young University and PM. Overall, DNA sequencing runs were of varying depths, and mitochondrial genomes were assembled using both reference-guided and de novo methods (Supplementary Methods & Results). We incorporated publicly available Rainbow Trout (*Oncorhynchus mykiss)* and Chinook Salmon (*Oncorhynchus tshawytscha*) mitogenomes in all downstream analyses. Rainbow Trout are the sister species to Cutthroat Trout (Smith & Stearley, 1989) and provide context to the branch lengths found between Cutthroat Trout subspecies.

Chinook Salmon represent a robust outgroup to root the trees as they are closely related to both Pacific Trout species (Behnke, 2002; Brunelli et al., 2013; Wilson & Turner, 2009).

### Mitochondrial Genome Annotation and Sequence Comparison

Each mitochondrial genome was annotated for all tRNA, rRNA, and protein-coding genes using MitoFish v4.09 (Iwasaki et al., 2013; Sato et al., 2018; Zhu et al., 2023). These genes were aligned using Mafft v7.525 (Katoh & Standley, 2013). A concatenated sequence super matrix was created using FASconCAT v1.7 (Kück & Longo, 2014). Mega v12 was used to calculate average pairwise percent differences between each sample in this concatenated sequence super matrix (Kumar et al., 2024), so that the genetic distance between Chinook Salmon, Rainbow Trout, and all the Cutthroat Trout lineages could be estimated.

### Phylogenetic Analyses

ModelFinder in IQtree v2.3.4 was run twice to find the best model of sequence evolution for each gene: once to test all models of sequence evolution available in IQtree v2.3.4, and again to test all models of sequence evolution available in BEAST v2.6 (Minh et al., 2020). IQtree v2.3.4 was used to generate a maximum likelihood phylogeny using 1000 ultrafast bootstrap alignments and 1000 iterations (Minh et al., 2020). Two Bayesian phylogenies were generated using BEAST v2.6 using the Yule process of speciation with a chain length of 90,000,000, one with a strict molecular clock, and one with an optimized relaxed molecular clock (Bouckaert et al., 2014).

Both Bayesian Trees were annotated using TreeAnnotator v2.7.6 using common ancestor heights, and a burn-in percentage of 10 percent (Drummond & Rambaut, 2007). Tracer v1.7.2 was used to assess adequate chain mixing by ensuring that all estimated sample sizes were over 200 for each parameter (Rambaut et al., 2018). We used treePL v1.0 to date our maximum likelihood phylogeny (Smith & O’Meara, 2012) and calibrated it using an *Oncorhynchus belli* fossil, setting the most recent common ancestor between all Rainbow Trout and Cutthroat Trout lineages between 9.7 and 10.7 Ma (Stearley & Smith, 2016). Cross-validation was used to choose the best smoothing value for the molecular clock.

## Results

Overall, the varying sequencing techniques and assembly methods all yielded viable mitochondrial genomes, leading to the inclusion of 148 mitochondrial genomes in this study (Supplementary Methods & Results). Annotated genomes were submitted to GenBank under BioProject accession number PRJNA1301035.

### Pairwise Sequence Divergence

The pairwise distance between all Cutthroat Trout lineages was very similar, with an average distance of 1.30% (Figure 2). Notably, the Bear River Cutthroat Trout and Yellowstone Cutthroat Trout had the smallest pairwise distance at 0.27%, while the largest sequence divergence was registered between Yellowfin Cutthroat Trout and Coastal Cutthroat Trout at 1.97%. The greatest pairwise distance observed in this entire sample set was between Chinook Salmon and the Yellowfin Cutthroat Trout at 7.19%, while all combined Rainbow Trout and Cutthroat Trout lineages differed by 4.98%.

**Figure 1.**
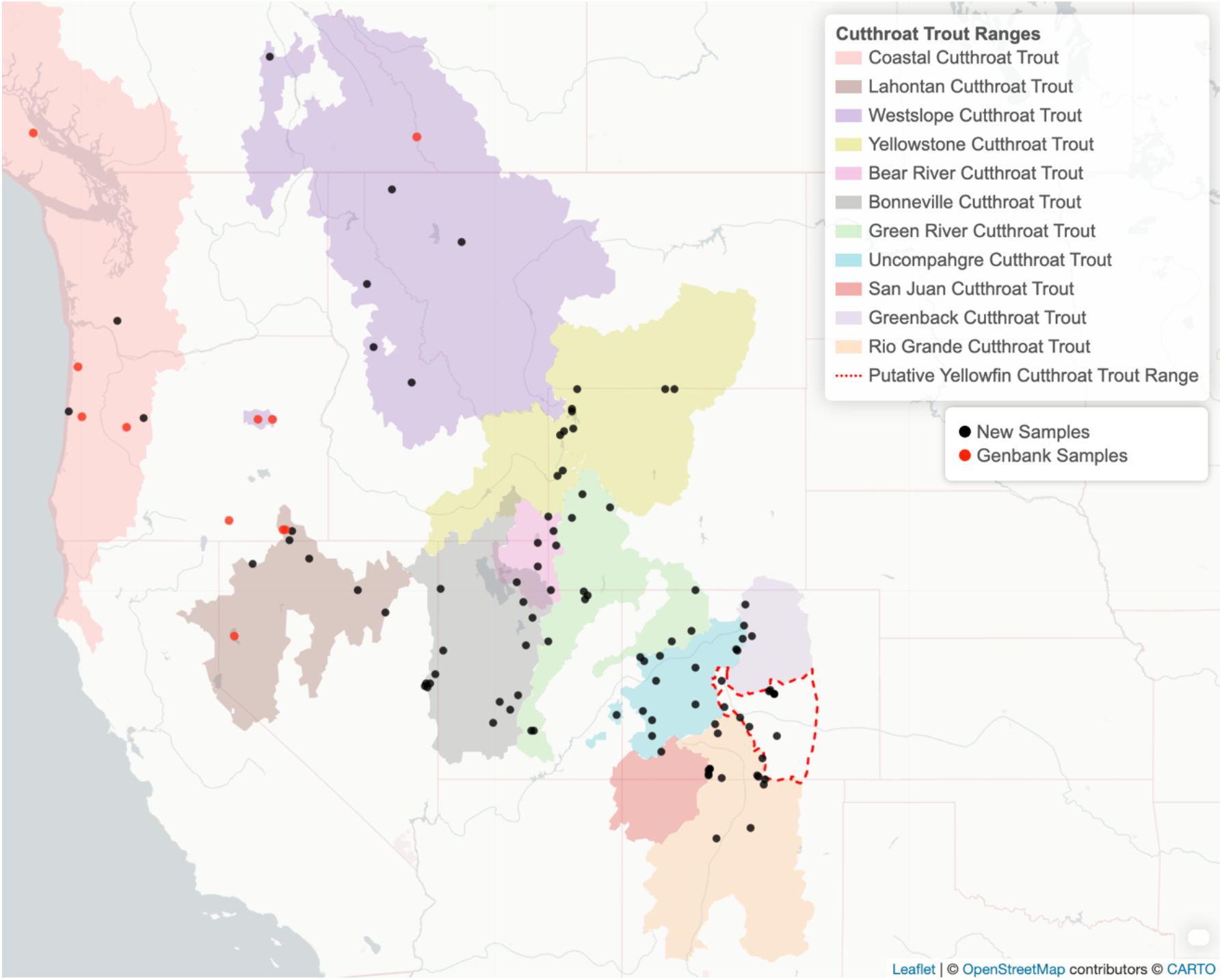
Native ranges of Cutthroat Trout in North America. Source locations for new samples acquired for this study are indicated with black circles, while red ones represent incorporated sequence data that was published in GenBank. Cutthroat Trout Native Ranges adapted from both the literature and www.nativetroutflyfishing.com.

**Figure 2.**
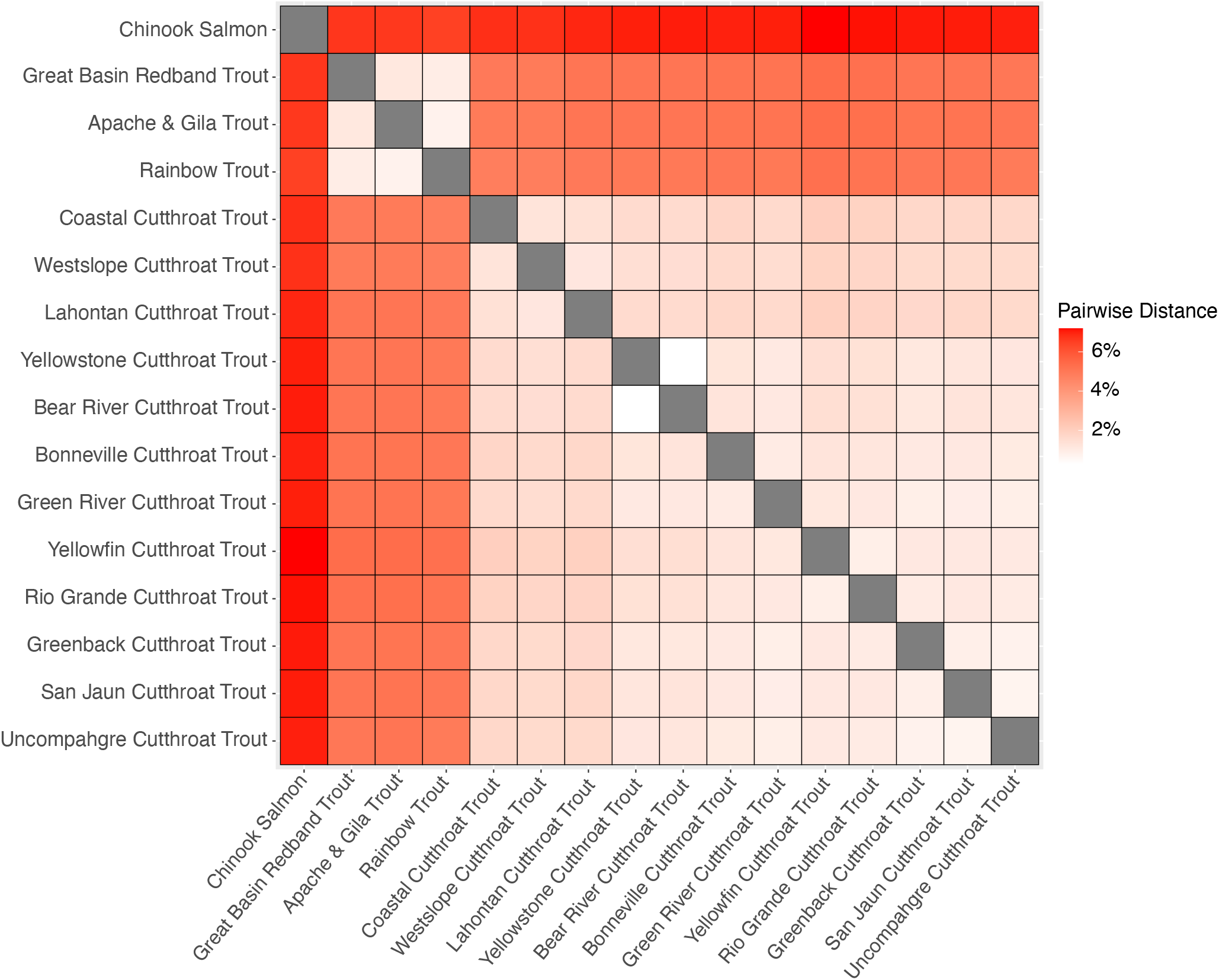
Pairwise average genetic distances between lineages of Cutthroat Trout, Rainbow Trout, and Chinook Salmon.

### Phylogenetic Analyses

The maximum likelihood phylogeny found twelve mitochondrial haplogroups of Cutthroat Trout separated by bootstrap values ranging from 77 to 100 (Figure 3 & Supplementary Figure 1). Both Bayesian phylogenies also found twelve mitochondrial haplogroups of Cutthroat Trout, with the strict molecular clock phylogeny having posterior probabilities that ranged from 0.85 to 1, and the optimized relaxed molecular clock having posterior probabilities that ranged from 0.71 to 1 (Figure 4 and Supplementary Figure 2). The strict molecular clock had a minimum effective sampling size of 1661, while the minimum effective sampling size of the optimized relaxed clock had a minimum effective sampling size of 208, indicating a high level of Markov Chain Monte Carlo chain convergence. TreePL showed that although Cutthroat Trout and Rainbow Trout diverged >10.2 million years ago, Coastal Cutthroat Trout did not diverge from other Cutthroat Trout lineages until >5 million years ago (Figure 3). Our molecular clock identified at least 11 lineages of Cutthroat Trout that diverged > 2 million years ago, and at least 13 lineages that diverged > 1 million years ago (Figure 3).

**Figure 3.**
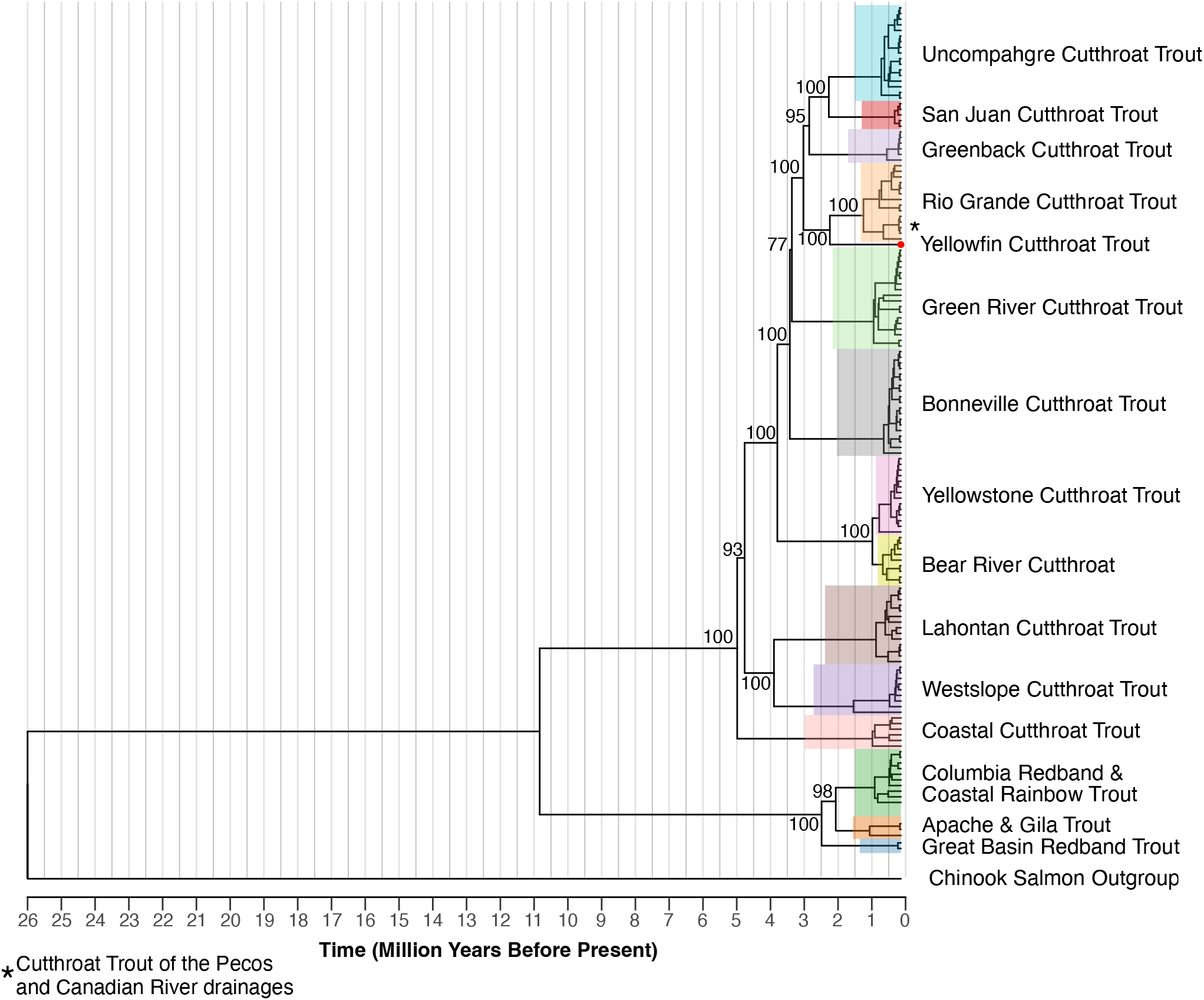
Dated maximum likelihood mitochondrial phylogeny with the most common ancestor of Cutthroat Trout and Rainbow Trout calibrated between 9.7 and 10.7 Ma. Bootstrap values (n = 1000) are shown for each node. The scale bar represents time in millions of years before the present.

**Figure 4.**
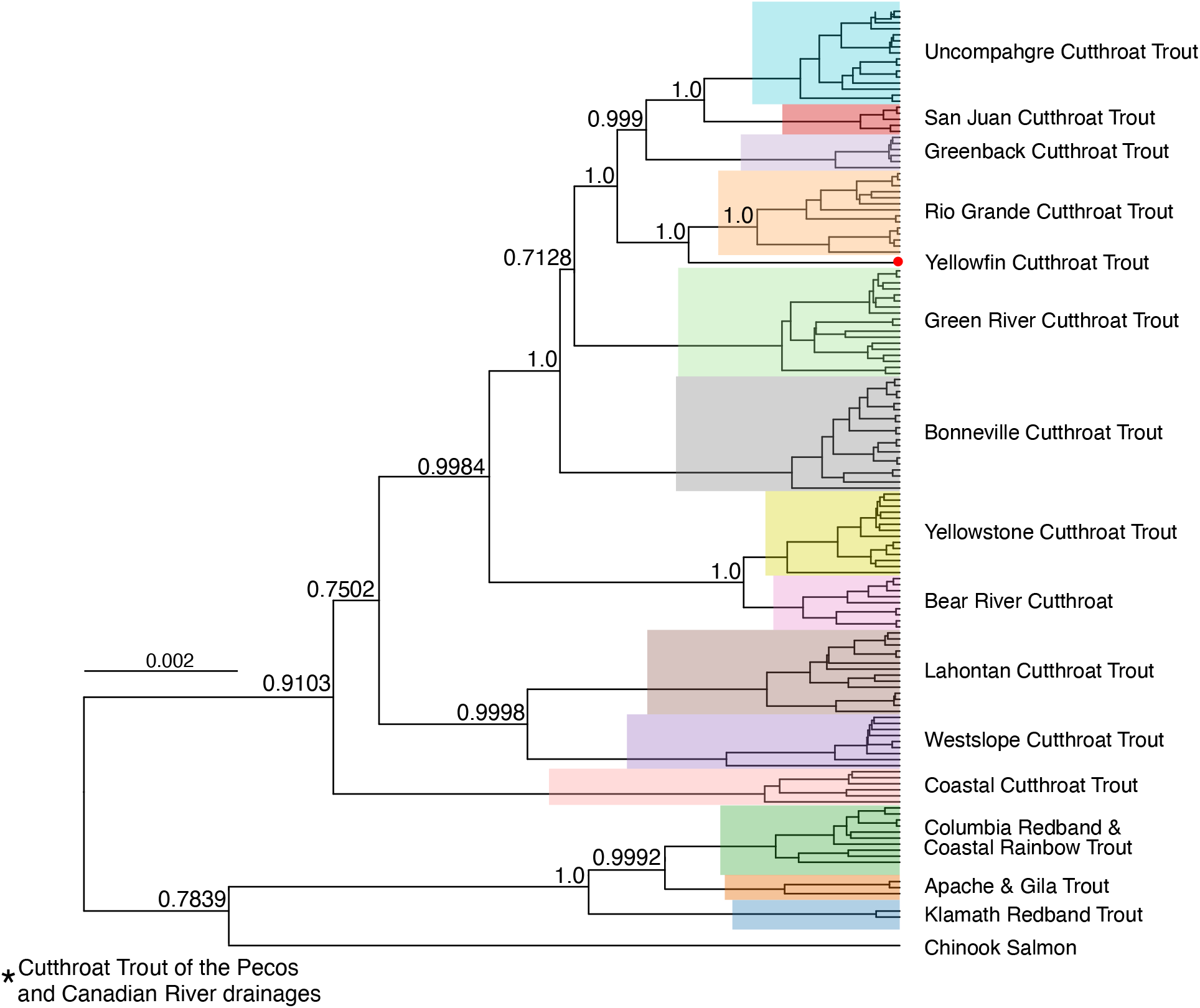
Optimized relaxed molecular clock Bayesian phylogeny of Cutthroat Trout mitochondrial genomes. Posterior probabilities are shown for each node.

## Discussion

Our phylogenies provide strong support for twelve mitochondrial haplogroups of Cutthroat Trout (Figure 3, 4, Supplementary Figure 1, 2). Excluding the Bear River – Yellowstone Cutthroat Trout haplogroup, all remaining pairwise distances ranged from .65% to 1.97%, highlighting a similar level of divergence between lineages (Figure 2). These findings are consistent with Behnke’s 1992 and 2002 phylogenies based on meristic counts, morphometric measurements, coloration, spotting patterns, and geographic distribution. Behnke (2002) recognized 14 subspecies of Cutthroat Trout, organized among four “major” subspecies: Coastal Cutthroat Trout (*O. c. clarkii*), Westslope Cutthroat Trout (*O. c. lewisi*), Lahontan Cutthroat Trout (*O. c. henshawi*), and the Yellowstone Cutthroat Trout complex (*O. c. bouvieri*). New fossil evidence shows that the ancestral Cutthroat Trout species (*Oncorhynchus belli*) had split from Rainbow Trout by at least 10.2 Ma (Stearley & Smith, 2016), highlighting that the split occurred earlier than Behnke hypothesized. However, despite the Rainbow-Cutthroat split having occurred by at least 10.2 Ma, our findings suggest that the diversification we see between different Cutthroat Trout lineages did not start until five million years ago (Figure 3). Overall, these analyses have provided more evidence to further refine the taxonomic relationships between different Cutthroat Trout lineages.

Phylogenetic relationships among Cutthroat Trout inferred in the past have used allozymes (Allendorf & Leary, 1988; Loudenslager & Gall, 1980; Utter & Allendorf, 1994; Shiozawa & Williams, 1992), partial mtDNA (Bestgen et al., 2019; Loxterman & Keeley, 2012; Metcalf et al., 2007, 2012; Pritchard et al., 2009; Shiozawa et al., 2018; Wilson & Turner, 2009) Y-chromosome sequences (Brunelli et al., 2013), microsatellites (Metcalf et al., 2007; Pritchard et al., 2007), Amplified Fragment Length Polymorphisms (Bestgen et al., 2019; Metcalf et al., 2007), and nuclear SNPs (Houston et al., 2012). Despite the broad array of molecular tools used to characterize phylogenetic relationships among Cutthroat Trout, the results have been surprisingly consistent and have revealed inaccuracies in Behnke’s taxonomy at the subspecies level. Additional studies with even higher resolution covering the nuclear genome continue to be produced. Kokkonen et al. (2024) examined 1827 nuclear genes and found many of the same relationships identified in this study. Clark (2025) also produced consistent phylogenies examining 40 million SNPs spread across the nuclear genome. Interestingly, her work revealed some discord between what we and others have found in the mitochondrial DNA compared to her nuclear phylogenies. For example, San Juan Cutthroat Trout (*O. c. ssp*..) nuclear DNA aligns more closely with Green River Cutthroat Trout (*O. c. pleuriticus*), while their mitogenomes are most closely related to Uncompahgre Cutthroat Trout (*O. c. ssp*..). In addition, the nuclear genome of South Hayden Creek fish (East Slope Uncompahgre Cutthroat Trout) aligns with Rio Grande Cutthroat Trout (*O. c. virginalis*), while the mitogenome is clearly Uncompahgre Cutthroat Trout. This discord reflects additional unexplained relationships between different Cutthroat Trout evolutionary lineages and highlights the need for additional work to continue to resolve the convoluted taxonomy of Cutthroat Trout. However, much can be gleaned from the full mitogenome phylogenies presented here – particularly with regard to some subspecies-specific findings that warrant further discussion:

### Coastal Cutthroat Trout

Behnke (2002) hypothesized that the ancestor of all Cutthroat Trout lived in a marine environment, and this is supported by the Coastal Cutthroat Trout as being the basal lineage using partial mitochondrial genomes (Bestgen et al., 2019; Loxterman & Keeley, 2012; Metcalf et al., 2007, Metcalf et al., 2012; Pritchard et al., 2009; Shiozawa et al., 2018; Wilson & Turner, 2009). Despite this, different data types, including allozymes and nuclear transcriptomes, have identified the Westslope Cutthroat Trout as being basal to all other subspecies of Cutthroat Trout (Allendorf & Leary, 1988; Kokkonen et al., 2024; Loudenslager & Gall, 1980; Utter & Allendorf, 1994; Shiozawa & Williams, 1992). With the allozyme and nuclear DNA phylogeny, natural Westslope Cutthroat Trout introgression with Rainbow Trout could also explain why phylogenetic analyses have placed the Westslope Cutthroat Trout as the most basal lineage.

While this study relies on mitochondrial DNA which does not show hybridization, two individuals identified as Westslope Cutthroat Trout were included in our study from the John Day River drainage, and both had Rainbow Trout mitochondrial genomes. This confirms that Westslope Cutthroat Trout-Rainbow Trout introgression exists in at least the John Day River System. Using 1827 nuclear genes to build a phylogeny of Cutthroat Trout, Kokkonen et al. (2024) showed that a Westslope Cutthroat Trout collected in Idaho’s Middle Fork Salmon River was the most basal Cutthroat Trout lineage. Westslope Cutthroat Trout have coexisted with Rainbow Trout for millennia (Behnke, 2002). Additionally, previous work has found widespread Rainbow Trout - Westslope Cutthroat Trout hybridization in the Salmon River System (Kozfkay et al., 2007). The high bootstrap (100) and posterior probabilities (1.0 with strict molecular clock, and 0.9103 with optimized relaxed clock) in our mitochondrial maximum likelihood study supports that the Coastal Cutthroat Trout is the most basal Cutthroat Trout lineage, and the misidentified John Day River Westslope Cutthroat Trout haplotypes suggest introgression may at least partially explain conflicting results from other studies.

### Lahontan Cutthroat Trout

One mitogenome of particular interest came from Guano Creek in Oregon. Despite having native Redband Trout (*O. m. ssp*..), Guano Creek has been stocked with multiple lineages of Cutthroat Trout, including Willow-Whitehorse Lahontan Cutthroat Trout, Heenan Lake Lahontan Cutthroat Trout, and Alvord Cutthroat Trout (*O. c. alvordensis*) (Behnke, 2007). Because of this, we expected that if a fish in Guano Creek had a Lahontan Cutthroat Trout mitogenome, it would be similar to the mitogenomes found in Willow-Whitehorse Cutthroat Trout, Heenan Lake Lahontan Cutthroat Trout, or the Alvord Cutthroat Trout. The Guano Creek mitogenome was identical to the haplotype found in Mahogany Creek, which drains into Summit Lake Nevada.

Importantly, one hypothesized entry point for Lahontan Cutthroat Trout into the Alvord Basin is via a headwater transfer of Mahogany Creek (Behnke, 2002; Curry & Melhorn, 1990). This could mean that this mitogenome may have come from the extinct Alvord Cutthroat Trout.

Additional comparisons need to be made to partial mitochondrial genomes from known formalin-fixed Alvord Cutthroat Trout, and other Lahontan Cutthroat Trout mitogenomes from Heenan Lake to confirm this hypothesis.

### Bonneville Cutthroat Trout

One needed revision to the taxonomy of Cutthroat Trout is the grouping of native trout in the Bear River Basin (Bear River Cutthroat Trout) with Bonneville Cutthroat Trout (*O. c. utah*) from the southern (main) part of the Bonneville Basin. Behnke hypothesized that the Bear River Cutthroat Trout gave rise to the Bonneville Cutthroat Trout when the Bear River was diverted into the Bonneville Basin approximately 30,000 years ago (Shiozawa et al., 2018), consistent with the hypothesis that Cutthroat Trout diversified ~1-2 Ma (Behnke, 2002; Behnke, 1992).

However, new fossil evidence suggests that divergence occurred much earlier at 10.2 MYA. Further, numerous genetic studies based on allozymes, mitochondrial DNA, and, recently, nuclear genes, have found that the Bear River Cutthroat Trout is sister to Yellowstone Cutthroat Trout, not the Bonneville Cutthroat Trout (Houston et al., 2012; Kokkonen et al., 2024; Loxterman & Keeley, 2012; Shiozawa & Evans, 1995; Toline et al., 1999). Results from these previous genetic studies and from this study suggest that the Bear River Cutthroat Trout should be considered a distinct lineage most closely allied with the Yellowstone Cutthroat Trout clade. While Loxterman & Keeley (2012) recommended that the Bear River Cutthroat Trout be called the Bonneville Cutthroat Trout, and those in the Southern (Main Basin) Bonneville Basin be called the Great Basin Cutthroat Trout, the type specimen for the Bonneville Cutthroat Trout is from Utah Lake, which is in the central part of the Main Bonneville Basin. Because of this, we recommend that Cutthroat Trout found in the Bear River drainage be called the Bear River Cutthroat Trout. We also recommend that the Bonneville Cutthroat Trout be recognized as having a distinct evolutionary history, separate from the Yellowstone Cutthroat Trout.

Additionally, our phylogeny shows that one Bear River Cutthroat Trout from Giraffe Creek in Wyoming falls within the Yellowstone lineage, and one Yellowstone Cutthroat Trout from Dog Creek (Teton River Tributary) Idaho, falls within the Bear River lineage, further complicating the relationship between Bear River Cutthroat Trout and Yellowstone Cutthroat Trout. While Yellowstone Cutthroat Trout derived from wild spawn operations in Yellowstone National Park were widely stocked across the Western United States, Giraffe Creek has a more basal haplotype that is not commonly found within Yellowstone National Park. This haplotype, along with the Bear River haplotype found in a Teton River tributary, could be explained by a historic connection between the Bear River and Snake River. A haplotype similar to the Bear River haplotype is also found in the middle Snake River of Idaho, suggesting that Bear River–like haplotypes were already unique prior to the diversion of the Bear River into Lake Bonneville.

This close relationship between Bear River Cutthroat Trout and Middle Snake River Yellowstone Cutthroat Trout could be a possible explanation for the Bear River haplotype found in Dog Creek. Despite these repeated connections, one Cutthroat Trout from Woodruff Reservoir in Utah was found to have a Yellowstone Cutthroat Trout haplogroup that matches what would be expected from Yellowstone Cutthroat Trout with an origin in Yellowstone National Park.

Even though there is a close genetic relationship shared between Yellowstone Cutthroat Trout and Bear River Cutthroat Trout, this Yellowstone Lineage haplotype is likely the result of stocking (Varley, 1980). More genetic analyses are needed to understand the range of native haplotypes found within both basins. Museum specimens collected prior to the advent of large-scale stocking could aid in this discovery if available (Metcalf et al. 2012).

### Cutthroat Trout of the Southern Rocky Mountains

The Southern Rocky Mountain (SRM) region historically included the Colorado River Cutthroat Trout (*O. c. pleuriticus*), Greenback Cutthroat Trout (*O. c. ssp*..), and Rio Grande Cutthroat Trout, along with the extinct Yellowfin Cutthroat Trout (*O. c. macdonaldi*; Behnke 1992). Mining and other anthropogenic pressures in the late 1800s resulted in native trout populations being over-exploited or eliminated, especially on the east slope of the Rocky Mountains. Introduction of nonnative trout across their native range resulted in further, more permanent declines to native trout populations (Adams, 1999; Behnke, 1992; McHugh & Budy, 2006; Peterson et al., 2013; Peterson et al., 2004; Shepard, 2004; Shepard et al., 2002). Using mostly meristic characters, Behnke (1992) found that the Greenback Cutthroat Trout, historically native to the South Platte and Arkansas River basins, were still represented by a few scattered populations in both river basins. A decades-long conservation effort used these fish to repopulate the headwaters of the South Platte and Arkansas basins with the goal of removing them from the federally threatened species list (AMEC, 2014). Work by Metcalf et al. (2007, Metcalf et al. 2012) determined that most of those relict populations were in fact lineages of Colorado River Cutthroat Trout. By the early 1900s, native trout from Trappers Lake in the headwaters of the White River (tributary of the Green River) were stocked widely across Colorado (Rogers et al., 2018a). In addition, fertilized Cutthroat Trout eggs were taken from the Grand Mesa in the headwaters of the Gunnison River basin in the late 1800s, and distributed into barren waters across the state, including the east-slope of the Rocky Mountains. This largely undocumented history of translocations led to long-term taxonomic confusion with extraordinary consequences (Metcalf et al. 2007, Metcalf et al. 2012; Bestgen et al. 2019). Since this discovery, the Greenback Cutthroat Trout recovery effort has been resurrected with the single remaining population native to the South Platte basin. Three lineages of Colorado River Cutthroat Trout are now recognized (Metcalf et al. 2012; Rogers et al. 2018a). These include 1) Green River Cutthroat Trout native to the Green, White, and Yampa river systems along with the Dirty Devil and Escalante rivers in Utah (formerly “blue” lineage sensu Metcalf et al. 2012; Rogers et al. 2018a; Bestgen et al. 2019), 2) Uncompahgre Cutthroat Trout native to the upper Colorado, Gunnison, and Dolores river basins (formerly “green” lineage sensu Metcalf et al. 2012; Rogers et al. 2018a; Bestgen et al. 2019), and 3) the San Juan Cutthroat Trout native to its namesake basin (Rogers et al., 2018b). These common names reflect the directed consensus of the Colorado River Cutthroat Trout Conservation Team charged with managing all three of these taxa (Rogers et al., 2025).

Despite this complicated history of stocking, over-exploitation, nonnative invasions, and taxonomic confusion, samples in this study reflect all of the major groups of Cutthroat Trout that would have been historically found in the SRM. Multiple ice ages have occurred since Cutthroat Trout invaded the SRM, leading to cyclic range expansions and contractions, eventually giving rise to the six Cutthroat Trout haplogroups we see today (Figures 3, 4). As such, we recognize the Green River Cutthroat Trout, Uncompahgre Cutthroat Trout, San Juan Cutthroat Trout, Greenback Cutthroat Trout, Yellowfin Cutthroat Trout, and the Rio Grande Cutthroat Trout as distinct lineages worthy of conservation.

Additionally, while not as distinct as the above lineages, Cutthroat Trout found in upper tributaries of both the Canadian and Pecos Rivers of New Mexico appear to have split from other Rio Grande Cutthroat Trout over one million years ago. Overall, the great diversity of Cutthroat Trout found in the SRM region highlights an ancient and complex evolutionary history.

### Major vs Minor Species Conflict

One of the most intractable challenges in taxonomy hinges on determining an appropriate threshold for delineating species, subspecies, and lineages. While seemingly academic, this problem has created many difficulties for species conservation and management. Inconsistent application across species (Mayden, 1999) has compounded the problem. Different taxa have varying characteristics that make determining this threshold difficult, including genetically distinct species with overlapping ranges (Cooke et al., 2012; Kautt et al., 2020; Lescroart et al., 2023; Stuart et al., 2006), morphological but not genetic differentiation (Cole et al., 2008; Miller & Benzie, 1997), genetic but not morphological differentiation (Cooke et al., 2012; Petzold & Hassanin, 2020; Stuart et al., 2006; Xue et al., 2021), huge geographic ranges with variable morphology (Joshi et al., 2023; Mutanen et al., 2012), and other challenges. Cutthroat Trout exhibit many of these confounding issues, including a large geographic range, lineages with morphological variation with little genetic differentiation (Lahontan and Paiute Cutthroat Trout & Snake River Cutthroat Trout), apparent sympatric ranges with genetic differentiation (Yellowfin and Uncompahgre Cutthroat Trout), and lineages with subtle morphological differences but substantial molecular ones (Uncompahgre Cutthroat Trout and Green River Cutthroat Trout). In part, these reasons suggested the need to consider revising the taxonomy of Cutthroat Trout, leading to a special workshop at the 2015 annual meeting of the American Fisheries Society to address the issue.

Since this workshop, Trotter et al (2018) proposed elevating the four major lineages described by Behnke as major subspecies to full species, which has been met with scrutiny. This argument was largely based on geographic, ecological, and morphometric data supporting the existence of these four groups, and perhaps a way to honor Behnke’s legacy since “major” subspecies are not recognized by the International Commission of Zoological Nomenclature. The four “major” subspecies described by Behnke were based on the assumption that Coastal, Westslope, Lahontan, and Yellowstone Cutthroat Trout split from each other one million years ago, with “minor” subspecies arising over the last 100,000 years (Behnke, 2002). Our calibrated phylogeny revises these dates, and our pairwise comparisons highlight a much deeper evolutionary history within Cutthroat Trout (Figures 3).

To add additional branch length context, we included the Rainbow Trout clade: Coastal Rainbow (*O. mykiss irideus*) and Columbia River Redband (*O. mykiss gairdnerii*), Apache (*O. apache*), Gila (*O. gilae*), and Great Basin Redband Trout (*O. mykiss newberii*). Our phylogenies indicate that Apache and Gila Trout are more closely related to the Rainbow Trout Clade than the Great Basin Redband Trout, yet Apache and Gila Trout are currently listed as distinct species, while Great Basin Redband Trout are considered a subspecies of Rainbow Trout under (*O. mykiss*). This taxonomic inconsistency reinforces the core challenge we face with Cutthroat Trout species designations. While some types of data may support more similarities between Great Basin Redband Trout and the Rainbow Trout than between Apache Trout, Gila Trout, and Rainbow Trout, the maternally inherited mitochondrial genomes suggest a different evolutionary history. Overall, while the Rainbow Trout complex is in need of careful genetic reexamination, the Unified Species Concept was created to allow species designation despite conflicting diagnosability, ecology, and genetic differentiation (De Queiroz, 2007). The flexibility that the Unified Species Concept allows is an important shift that may make both Rainbow Trout and Cutthroat Trout taxonomy more straightforward.

The Unified Species Concept was created because, despite almost a century of debate, biologists have not agreed on what constitutes a species. Many of the species concepts that exist contradict each other and make species designation both more complicated and less consistent. The Unified Species Concept states that a species is a “separately evolving” metapopulation. While reproductive isolation, diagnosability, distinct ecologies, or monophyly can be evidence of differing evolutionary trajectories between metapopulations, these properties are no longer required. Additionally, De Queiroz (2007) makes it clear that the lack of any one property (reproductive isolation, diagnosability, distinct ecologies, monophyly, etc.) does not act as evidence contradicting a hypothesis of separate evolutionary trajectories, though more evidence does strengthen the hypothesis. This flexibility is the very reason that the Unified Species Concept was selected by Cutthroat Trout special workshop participants at the 2015 annual meeting of the American Fisheries Society. The ability to make species designations based on the best available data, while also allowing hypotheses to be strengthened and changed over time in response to newly collected evidence, is a species concept that does work for Cutthroat Trout taxonomy.

While this study does not evaluate reproductive isolation, diagnosability, or distinct ecologies, maternally inherited mitochondrial DNA supports the existence of twelve distinct lineages of Cutthroat Trout (Table 1). Furthermore, despite the Yellowfin Cutthroat Trout having been considered a “minor subspecies” by Behnke, the pairwise distance found between Yellowfin Cutthroat Trout and Yellowstone Cutthroat Trout is 1.42%, which is similar to the distances found between Yellowstone Cutthroat Trout and Lahontan, Westslope, and Coastal Cutthroat Trout (1.55%, 1.40%, and 1.53% respectively) highlighting that interim classification does not accurately describe the diversity found within Cutthroat Trout. In addition to the support found in our study, many previous studies have found these lineages consistently monophyletic across both mitochondrial and nuclear genomes (Houston et al., 2012; Kokkonen et al., 2024; Metcalf et al., 2007, Metcalf et al., 2012; Rogers et al., 2018a; Shiozawa et al., 2018) and morphologically distinct (Behnke, 2002, Behnke, 1992; Bestgen et al., 2013, 2019). Being both morphologically differentiated and monophyletic are characteristics that support these 12 lineages as being distinct but does not mandate discarding forty years of precedent. We see large differences between Rainbow Trout and Cutthroat Trout (species), and much smaller differences between these 12 lineages of Cutthroat Trout – consistent with the long-held practice of referring to them as subspecies. While other researchers have proposed splitting Cutthroat Trout into four species, our phylogeny does not suggest Coastal, Westslope, Lahontan, and Yellowstone Cutthroat Trout are substantially more distinct than the other eight lineages. As such, we continue to recognize a single Cutthroat Trout species (*Oncorhynchus clarkii*) here, comprised of twelve distinct subspecies (Table 1).

**Table 1.**
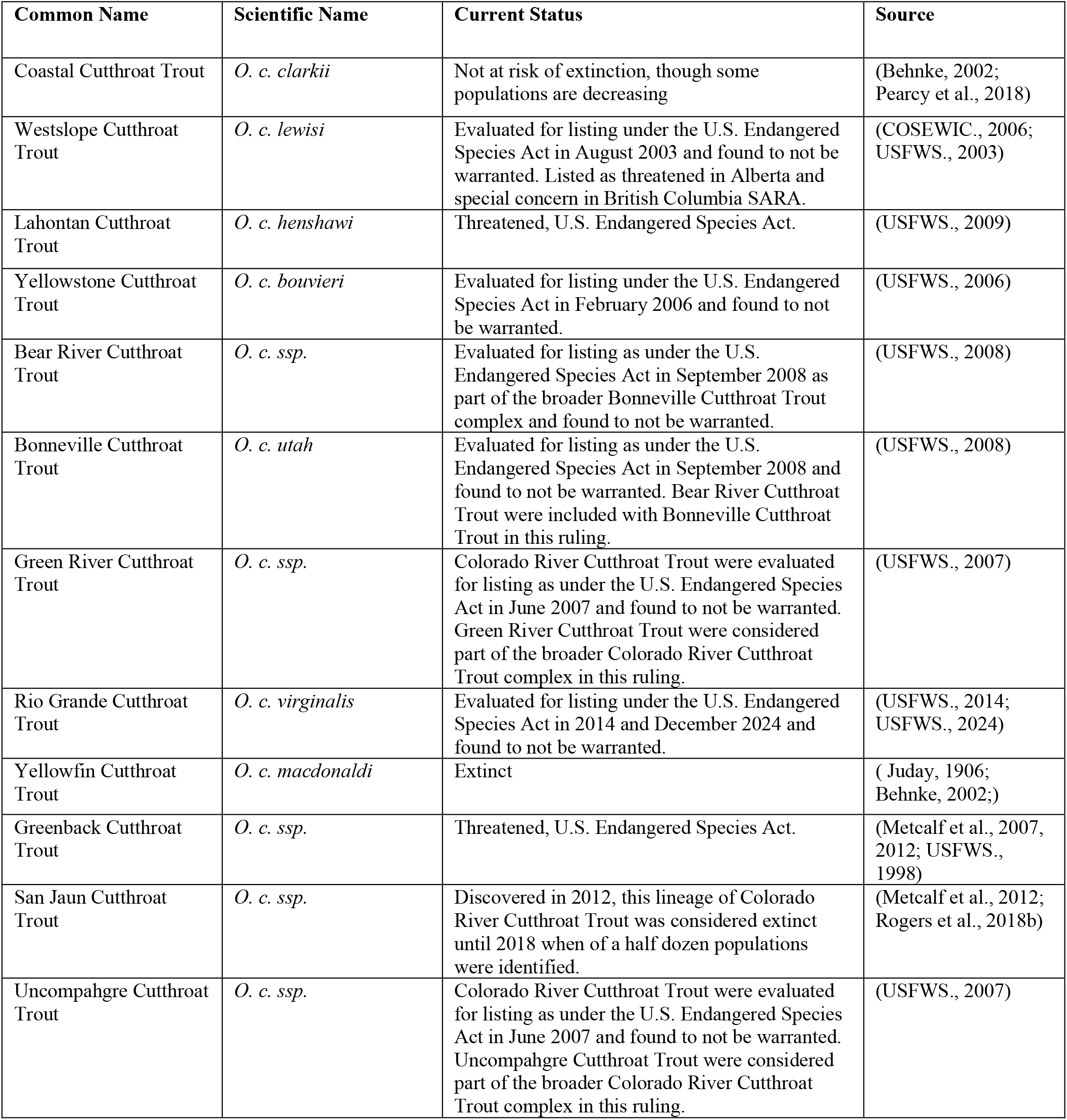
Mitochondrially distinct Cutthroat Trout lineages, including status assigned by the U.S. Fish and Wildlife Service (USFWS), Committee on the Status of Endangered Wildlife in Canada (COSEWIC), or other sources. Adapted from (Penaluna et al., 2016).

This arrangement acknowledges the profound diversity within Cutthroat Trout, aligns with the Unified Species Concept, facilitates consistency with existing conservation efforts, management plans, and ESA-related decisions, all while preserving stability associated with historic nomenclature. We acknowledge that continued research into the nuclear genome of Cutthroat Trout may provide evidence to further refine the taxonomic relationships between distinct lineages, but taxonomic changes should only follow if existing nomenclature is proven inaccurate.

### Conclusion

While additional work integrating calibrated phylogenies with geologic history will no doubt shed more light on possible dispersal routes across western North America and the Rocky Mountains, this work makes clear that Cutthroat Trout are a highly diverse and complex species, with divisions that are pronounced and ancient. The diversification between different Cutthroat Trout lineages started > 5 million years ago, with 11 lineages of Cutthroat Trout that diverged > 2 million years ago, and at least 13 lineages that diverged > 1 million years ago. Long branch lengths between haplogroups highlight their distinctiveness and help provide clarity for subspecies designations that have been contested for decades. Continuing to group all lineages as a single species under *Oncorhynchus clarkii* acknowledges their shared evolutionary history, allows consistency with the past 150 years of taxonomic discovery, and provides a framework for formal redescription of subspecies - critical for facilitating management and conservation activities for the myriad pieces that comprise Cutthroat Trout diversity.

## Supporting information

Supplementary Methods & Results

## Acknowledgments

We would like to thank the BYU Office of Research Computing for granting us access to computing power and for offering technical support. We would also like to thank Colorado Parks and Wildlife and the Brigham Young University Department of Microbiology and Molecular Biology for funding this project. Additionally, we thank the diligent team at Pisces Molecular in Boulder, Colorado for sequencing a large number of mitogenomes used in this study. We would also like to thank Gary Marsten for providing Cutthroat Trout range files and information on the Alvord Cutthroat Trout.

## Data Availability

The code underlying the genetic analyses in this article is available on Github at the link [https://github.com/Tsvanorden/Cutthroat_Mitochondrial_Genomes]. All mitochondrial genomes used in this study are publicly available under GenBank BioProject accession number PRJNA1301035

