## Supplementary Methods & Results for "Full Mitochondrial Phylogeny of Cutthroat Trout with Comments on Species Delimitation and Taxonomy"

**Supplementary Methods and Results:**

### Monte L. Bean Museum

*Sequencing and Mitochondrial Genome Assembly*

In all cases, whole genomic DNA was isolated using DNeasy® tissue kits (Qiagen, Germantown, MD, USA) following the manufacturer's recommended protocol. DNA purity and quantity was verified using a NanoDrop One (Thermo Fisher Scientific, Waltham, MA, USA) and a Qubit fluorometer with a dsDNA Broad Range assay kit. 85 DNA samples from MLBM were sent to Novogene America (Davis, California, USA), who prepared the libraries using the Illumina Nextera DNA Flex kit and sequenced the libraries with an Illumina Hi-Seq 2500 (Illumina, San Diego, CA, USA) using paired-end reads (2 x 150 bp) (Supplementary Table 1). DNA libraries were randomly distributed across 4 lanes, and sequencing was continued until each sample had yielded at least 2 GB of paired-end reads. The additional five DNA samples were sent to the University of Oregon Sequencing Center for library preparation and sequencing (Supplementary Table 1). Each library was verified using a MiSeq Nano and then sequenced on an Illumina NovaSeq - S4 - PE 150 Cycle (Illumina, San Diego, CA, USA) using paired-end reads (2 x 150 bp) for a minimum of 33 million paired-end reads per sample (16.5 million reads per direction). Raw reads from MLBM samples were assembled into full mitochondrial genomes using MitoZ v3.6 with megahit and kmer lengths of 59, 79, 99, 119, and 141 (Meng et al., 2019). As part of the innate MitoZ pipeline, fastp was used to trim out adapters and low-quality reads (Chen et al., 2018). For samples sequenced at Novogene America (Davis, California, USA), mitochondrial genome coverage ranged from 17.4x to 404.2x with an average of 58.5x coverage (Supplementary Table 1). For samples sequenced at the University of Oregon Sequencing Center, mitochondrial genome coverage ranged from 33.5x to 227.3x with an average of 179.7x coverage (Supplementary Table 1).

### Pisces Molecular

*Sequencing and Mitochondrial Genome Assembly*

Total genomic DNA was isolated using DNeasy tissue kits (Qiagen, Germantown, MD, USA) using the manufacturers recommended "mouse tail" protocol. Total mitochondrial DNA was amplified from each sample in three fragments, using primer pairs as follows: Fragment 1: forward primer in tRNA-Glx, reverse primer in ND3, 5975 bp. Fragment 2: forward primer in tRNA-Gly, reverse primer in CytB, 5972 bp. Fragment 3: Forward primer in CytB, reverse primer in ND2, 5635 bp. In the first round three amplified fragments for each of 8 samples were purified with Qiaquick PCR purification columns (Qiagen, Germantown, MD, USA) according to manufacturer's instructions, then sent to the University of Colorada BioFrontiers Institute for library preparation and sequencing (Supplementary Table 2). Libraries were prepared by tagmentation and indexed using an Illumina Nextera XT DNA Library Preparation kit, then sequenced with an Illumina MiSeq (Illumina, San Diego, CA, USA) using paired end reads (2 x 150 bp). In the second round, an additional 24 samples were prepared and sequenced at the BioFrontiers Institute with the same procedures (Supplementary Table 2). In the third round,

DNA was extracted and the three mitochondrial fragments PCR amplified and purified from one museum samples at the University of Colorado in a facility that had not previously been used for Cutthroat Trout samples (CU Cooperative Institute for Research in Environmental Sciences building), then sequencing carried out at the BioFrontiers Institute. Raw reads were processed to trim adapter and primer sequences then assembled into a full mitochondrial genome using Geneious R6.1 (Biomatters, Auckland, NZ). The mitochondrial genome from the historic Yellowfin Cutthroat Trout sample had an average of 14.1x coverage, with 100% being high-quality calls (Supplementary Table 2).
