## Supplementary Figures for "Full Mitochondrial Phylogeny of Cutthroat Trout with Comments on Species Delimitation and Taxonomy"

Supplementary Figure 1.

Undated maximum likelihood mitochondrial phylogeny. Bootstrap values (n = 1000) are shown for each node. The scale bar represents time in millions of years before the present.

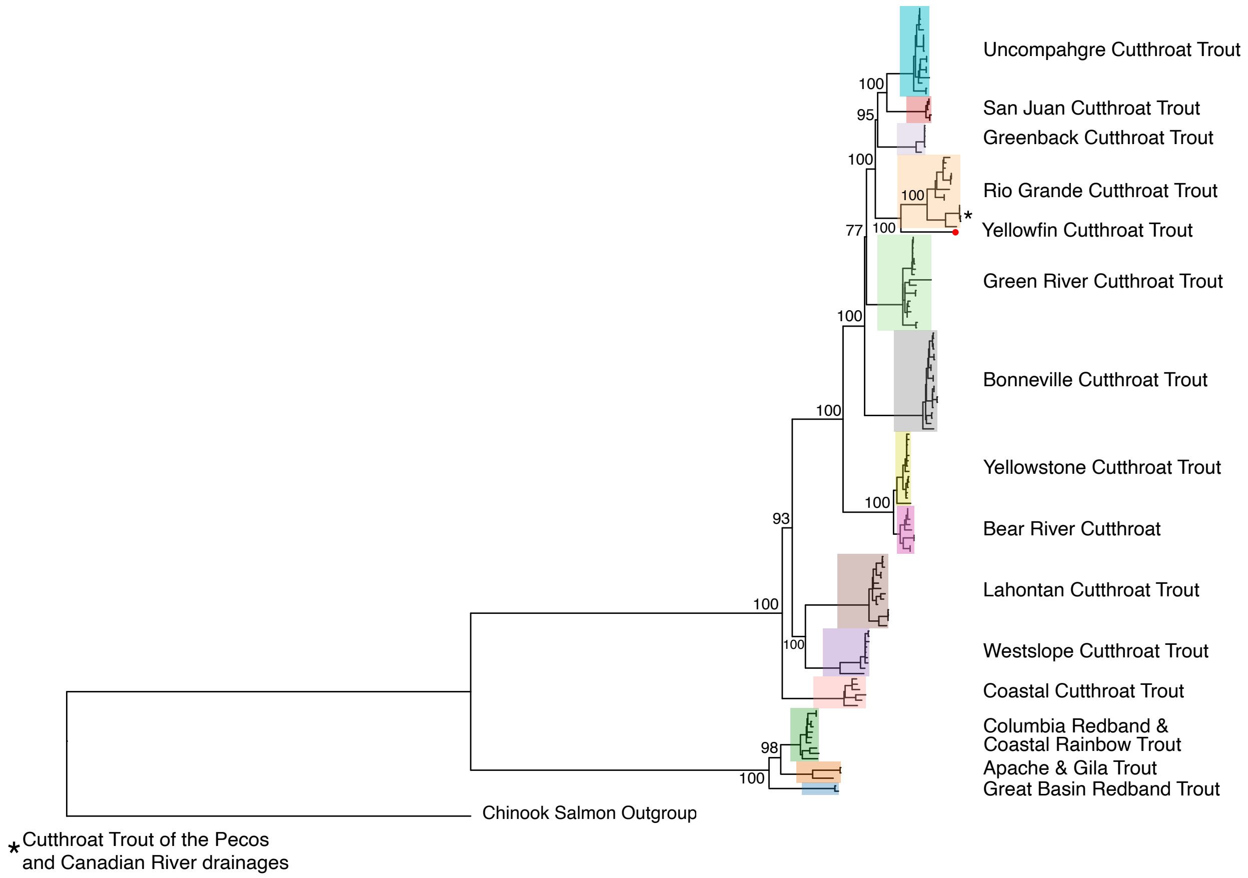

Supplementary Figure 2.

Strict molecular clock Bayesian phylogeny of Cutthroat Trout mitochondrial genomes. Posterior probabilities are shown for each node.

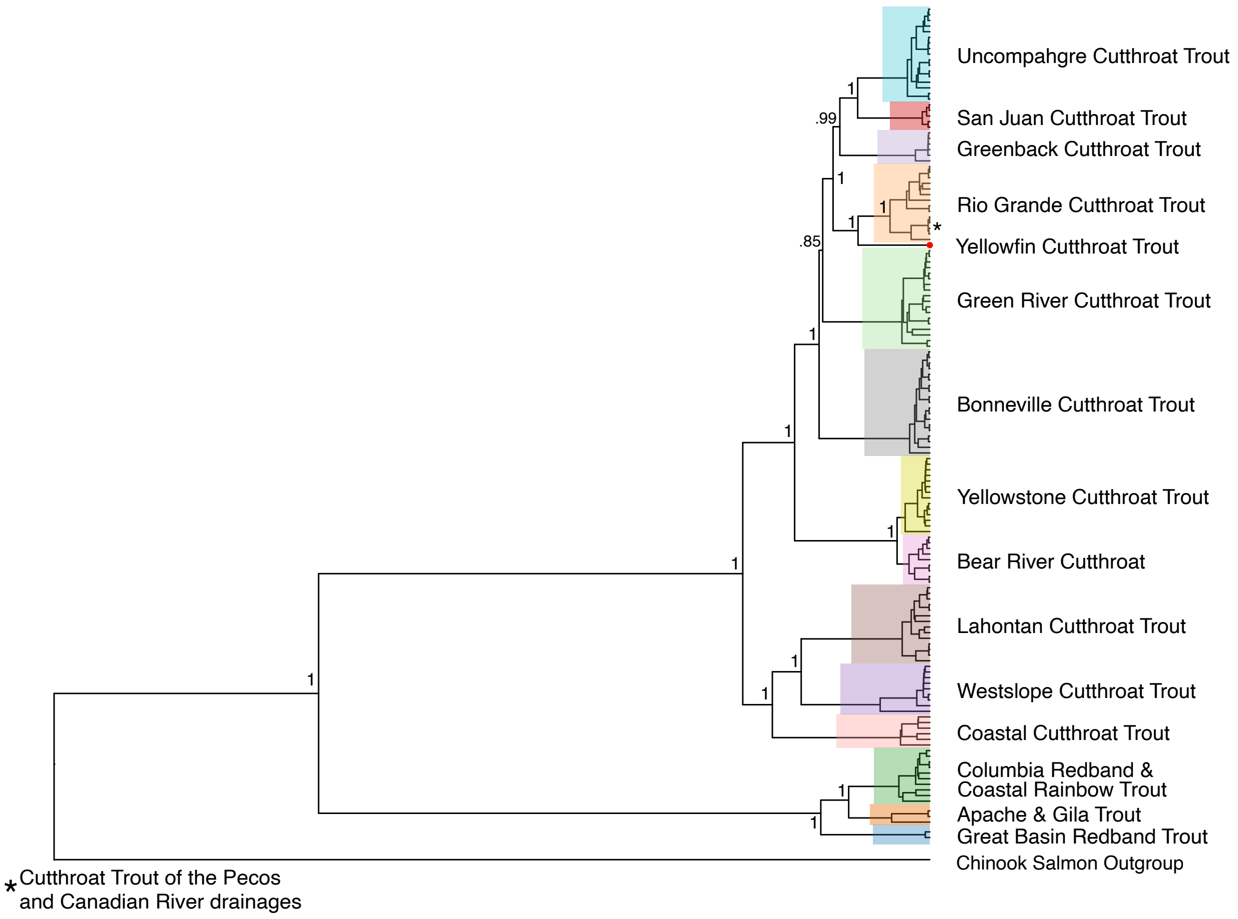
